# Optimizing 3D Spheroid Formation in Microwells via a Simple Pluronic F127 Coating

**DOI:** 10.64898/2026.08.18.744263

**Authors:** Nia Ho, Hiroyuki Kato, Hirotake Komatsu

**Affiliations:** Biophysics Program, Department of Natural Sciences, Pitzer College, 1050 N. Mills Ave., Claremont, CA 91711, USA; Division of Transplant Surgery, Department of Surgery, University of California, San Francisco, 513 Parnassus Ave., San Francisco, CA 94143, USA

**Author notes:** Corresponding author: Hirotake Komatsu.

**Keywords:** 3D cell culture, Pluronic F127, Microwells, Cell spheroids, Anti-adhesive surface coating, Surface modification, Tissue engineering, Regenerative medicine, High-throughput screening

## Abstract

Three-dimensional (3D) spheroid culture provides a physiologically relevant alternative to conventional two-dimensional culture, but reliable spheroid formation in microwells depends on limiting cell–substrate adhesion. Pluronic F127 is an amphiphilic triblock copolymer that forms a hydrated surface layer, reducing protein adsorption. Here, we evaluated whether this intrinsic anti-fouling property could restore an anti-adhesive surface in used microwell plates to promote spheroid formation. Using chondrogenic ATDC5 and pancreatic β-cell INS-1 cells, we characterized spheroid assembly kinetics, F127 cytotoxicity, surface hydrophilicity, protein adsorption, and spheroid morphology including size and shape factor. Both cell types formed compact spheroids within 24 hours on commercial anti-adhesive microwells. F127 coating markedly reduced water contact angle and protein adsorption, confirming increased surface hydrophilicity and reduced protein fouling. In microwells stripped of their original surface coating, F127 coating amounts of approximately 0.011–0.045 mg/cm^2^ consistently promoted spheroid formation across both cell types. Soluble F127 concentrations were confirmed to be non-cytotoxic up to 0.625% (w/v), while even complete dissolution of the highest tested coating amount would correspond to only 0.025% (w/v) F127. This simple, reproducible, and low-cost surface-modification strategy may provide an accessible approach for re-functionalizing microwell platforms for 3D cell culture.

## Introduction

Conventional biological testing heavily relies on flat, two-dimensional (2D) cell cultures that fail to replicate the complex architecture of three-dimensional (3D) human tissues. This structural limitation alters cellular morphology, surface receptor spatial organization, and downstream drug sensitivity^1,2^. To bridge this physiological gap, 3D multicellular spheroids have emerged as a robust platform that closely mimics native tissue geometry, cell–cell interactions, and in vivo-like gene expression profiles ^3^. Various methodologies exist to generate 3D spheroids, each presenting distinct technical trade-offs. While hanging-drop cultures follow a simple approach, they suffer from limited throughput and operational scalability. Conversely, dynamic bioreactors enable large-scale spheroid production with efficient nutrient supply, yet expose sensitive cells to damaging hydrodynamic shear stress and demand specialized equipment ^4^. Microwell array platforms represent another established approach, combining high-throughput capacity with precise control over aggregate size uniformity^5,6^. However, successful spheroid formation in microwells largely depends on the durability of an anti-adhesive surface layer that prevents cell– substrate attachment and promotes cell–cell aggregation ^7^.

Commercial low-attachment microwell plates typically rely on anti-fouling surface treatments that create a highly hydrophilic interface, thereby limiting protein adsorption and subsequent cell–substrate interactions. This functionally anti-adhesive surface favors cell–cell interactions and promotes spheroid assembly. However, many polystyrene microwell surfaces are intrinsically cell adhesive unless specifically modified, and low-attachment surface properties may also be diminished or lost during use. Pluronic F127 is a biocompatible amphiphilic triblock copolymer composed of poly(ethylene oxide)-poly(propylene oxide)-poly(ethylene oxide) (PEO-PPO-PEO) that forms a highly hydrated hydrophilic layer at aqueous interfaces ^8,9^. Its anti- fouling properties reduce protein adsorption and consequently limit cell–substrate adhesion ^10^. Therefore, surface functionalization with F127 may provide a simple and reproducible strategy for imparting anti-adhesive properties to microwell surfaces, including native polystyrene surfaces or surfaces that have lost their original low-attachment functionality, thereby promoting consistent 3D spheroid formation.

In this study, we systematically evaluated Pluronic F127 as a surface-modification agent to restore the anti-adhesive properties of microwell plates after primary culture use. We aimed to establish an optimized coating range that consistently produces round, uniform spheroids across multiple cell lines while maintaining cellular health and viability. Re-functionalizing used microwell platforms through this simple coating strategy provides a cost-effective and sustainable alternative to single-use commercial consumables ^2^. By reducing economic and resource barriers associated with high-throughput 3D culture, this accessible platform facilitates drug screening, regenerative medicine, and broader biological research ^2,11^, ultimately supporting the transition from conventional 2D testing to physiologically relevant biomimetic models.

## Results

### Pluronic F127 coating converted the polystyrene surface to a highly hydrophilic state

To establish the physical surface modifications imparted by Pluronic F127, we evaluated static water contact angles across a range of coating densities (0–0.0884 mg/cm^2^). Because degraded commercial microwell arrays expose underlying, untreated polystyrene, flat polystyrene substrates were used as a model interface to enable precise optical contact angle measurements without geometrical optical interference from array wells (**Fig. 1A**). Application of Pluronic F127 markedly enhanced surface wettability, as evidenced by a dramatic drop in static water contact angle compared with uncoated control substrates (0 mg/cm^2^; **Fig. 1A, B**). At low coating densities (< 0.0027), the contact angle decreased steeply from the native hydrophobic regime (> 60°) to < 15° (**Fig. 1B**). Beyond a coating threshold of 0.0027 mg/cm^2^, the surface reached a stable hydrophilic plateau (< 10°), with higher coating amounts up to 0.0884 mg/cm^2^ exhibiting no statistically significant differences in contact angle (*p* > 0.05; **Fig. 1B**). These physical characterization data demonstrate that Pluronic F127 adsorption rapidly converts native polystyrene into a highly, robust hydrophilic interface. Detailed statistical parameters are provided in **Supplementary Table 1**.

**Figure 1.**
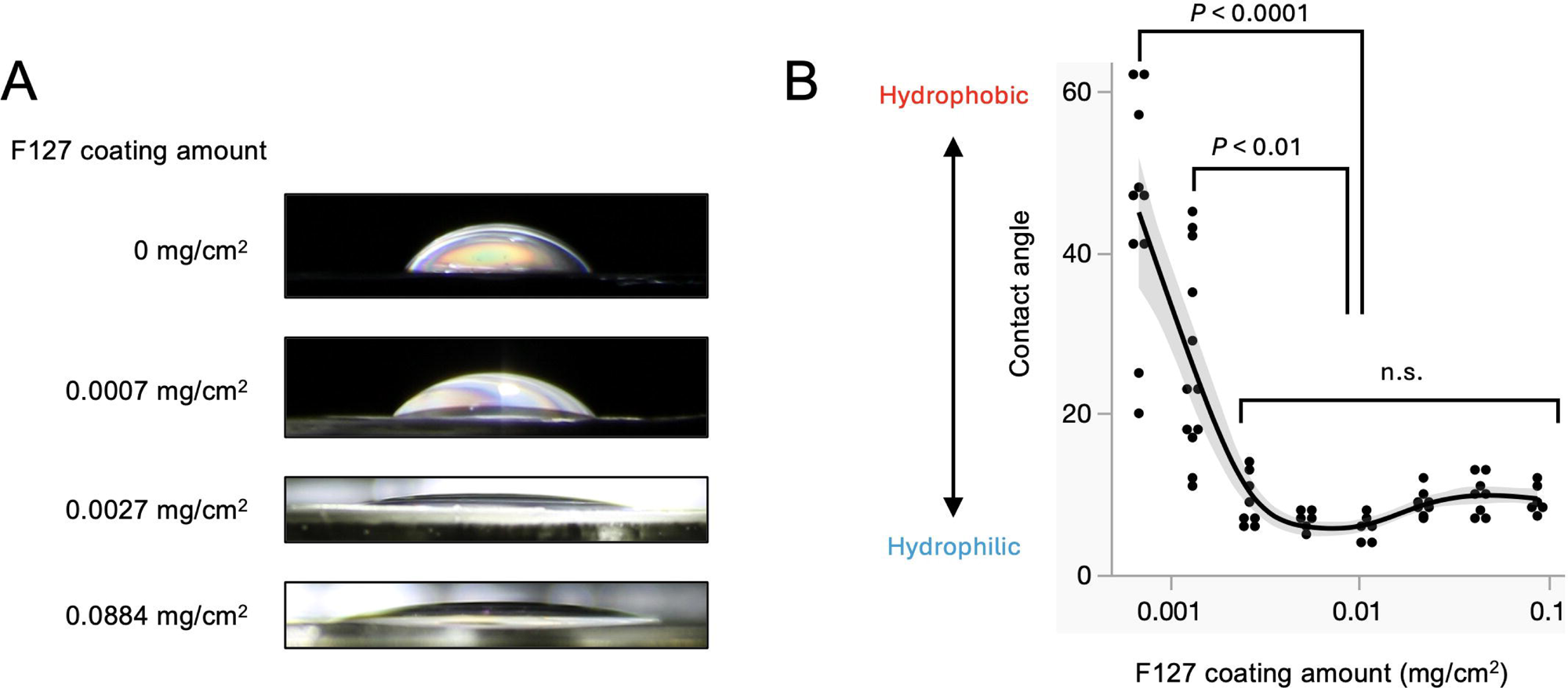
Surface hydrophilicity of Pluronic F127-coated polystyrene assessed by static water contact angle measurement. (A) Representative side-view images of 10 µL water droplets placed on flat polystyrene surfaces coated with Pluronic F127 at 0, 0.0007, 0.0027, or 0.0884 mg/cm^2^. (B) Quantification of static water contact angle across Pluronic F127 coating amounts. The solid line represents a smoothing spline fit (λ = 0.500), and the gray shaded area represents the confidence interval of the fitted curve. Individual points represent individual measurements. n = 6–12 per coating amount. Group differences were evaluated by one-way ANOVA with post hoc multiple comparisons (Tukey- Kramer HSD). n.s., not significant.

### Pluronic F127 coating reduced protein adsorption on stripped microwell surfaces

Because contact angle analysis confirmed that Pluronic F127 significantly enhances surface hydrophilicity–a key hallmark of an anti-adhesive hydration layer–we next evaluated its functional capacity to block protein adsorption using fluorescein isothiocyanate-labeled bovine serum albumin (BSA-FITC). We first stripped the original commercial low-attachment coating from microwell plates using 100% ethanol to expose the raw substrate and then re-coated the wells with designated amounts of Pluronic F127 (0 to 0.089 mg/cm^2^). We incubated the plates with BSA-FITC for 1 hour, washed them thoroughly to remove unbound protein, and measured the retained surface fluorescence (**Fig. 2**). The uncoated stripped microwells exhibited high baseline BSA-FITC adsorption (> 5.5 × 10^4^ RFU), whereas Pluronic F127 coating dramatically suppressed protein retention (**Fig. 2**). All tested coating densities (0.011–0.089 mg/cm^2^) significantly reduced fluorescence compared with the uncoated control (p < 0.0001). Furthermore, we observed no statistically significant differences in residual fluorescence among the F127-coated groups across this concentration range (p > 0.05). These results demonstrate that Pluronic F127 coating creates a robust anti-fouling barrier that effectively prevents non-specific protein adsorption on stripped microwell surfaces. Detailed statistical parameters are provided in **Supplementary Table 2**.

**Figure 2.**
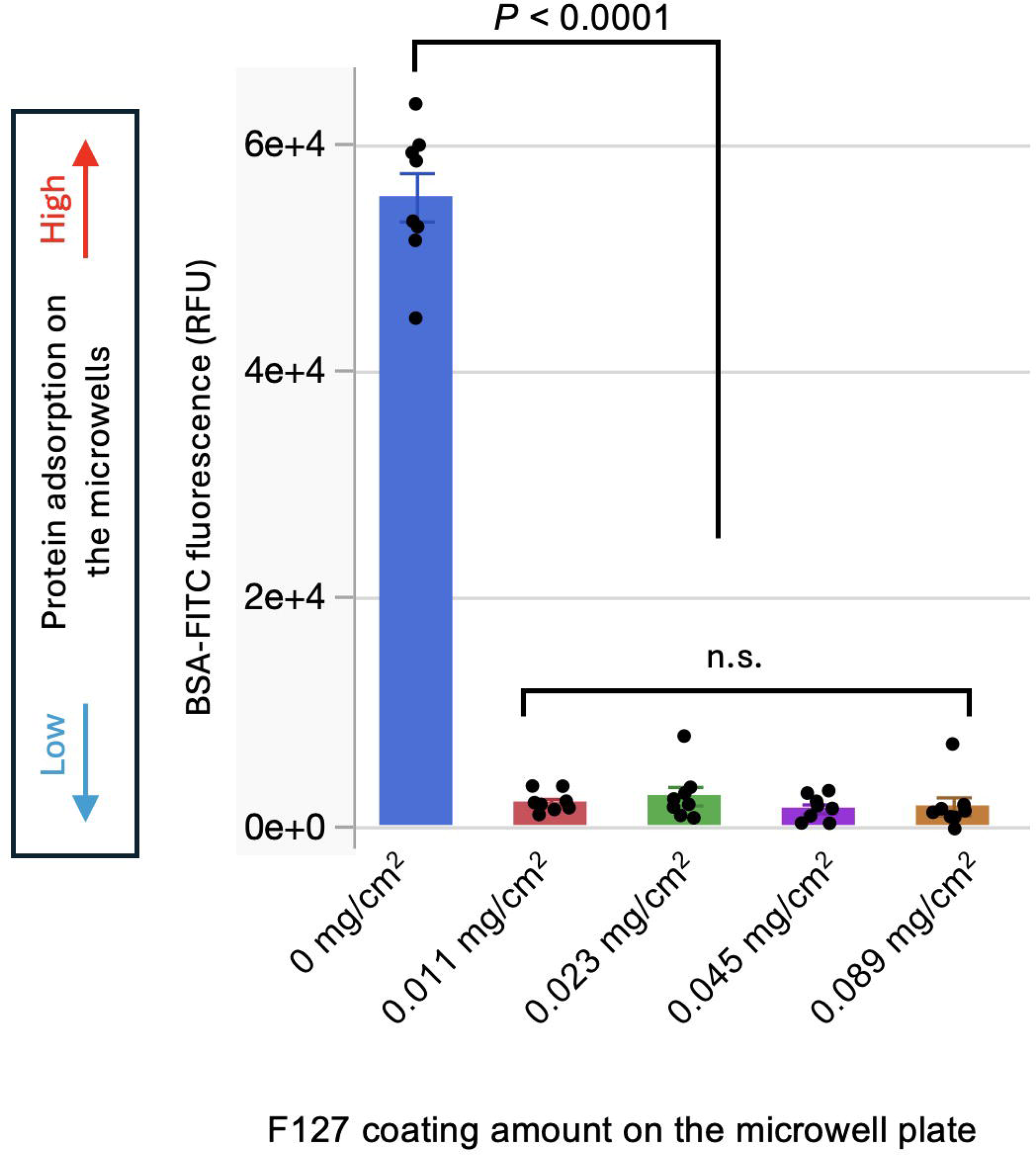
Protein adsorption on Pluronic F127-coated microwell surfaces assessed using BSA-FITC. Quantification of background-subtracted BSA-FITC fluorescence intensity (RFU) on uncoated and F127-coated microwell surfaces at coating amounts of 0, 0.011, 0.023, 0.045, and 0.089 mg/cm^2^. Individual points represent independent measurements, and bars represent mean ± SEM. n = 8 per condition. Group differences were evaluated by one-way ANOVA with post hoc multiple comparisons (Tukey-Kramer HSD). P < 0.0001 and n.s., not significant, are indicated in the figure.

### Low concentrations of Pluronic F127 in the culture medium preserved spheroid viability and morphology

We evaluated the cytotoxicity of soluble Pluronic F127 by exposing preformed ATDC5 and INS-1 spheroids to F127 concentrations ranging from 0% to 10% (w/v) for 24 h and quantifying spheroid viability and shape factor. Representative live/dead images showed predominantly viable spheroids at lower F127 concentrations, whereas higher concentrations progressively increased PI-positive non-viable regions (**Fig. 3A**). This concentration-dependent loss of viability appeared more prominently in INS-1 spheroids, particularly at 5% and 10% F127. Quantitative analysis showed that ATDC5 spheroids maintained relatively high viability across the tested concentrations, with only a modest decline at higher F127 concentrations (**Fig. 3B**). INS-1 spheroids showed greater sensitivity to soluble F127. Their viability remained relatively stable at lower concentrations but declined substantially at 5% and decreased markedly at 10%. Increasing F127 concentrations also altered spheroid morphology (**Fig. 3C**). ATDC5 spheroids showed only a modest reduction in shape factor across the tested range, indicating relatively preserved spheroid morphology. In contrast, INS-1 spheroids showed a progressive decrease in shape factor as the F127 concentration increased, with the largest disruption at 10%. Both cell types maintained shape factor without a significant decrease from the 0% control through 0.625% F127. Together, these results identify 0%–0.625% (w/v) F127 as a concentration range that preserves both viability and spheroid morphology across ATDC5 and INS-1 cells. **Supplementary Table 3** provides the detailed statistical analyses.

**Figure 3.**
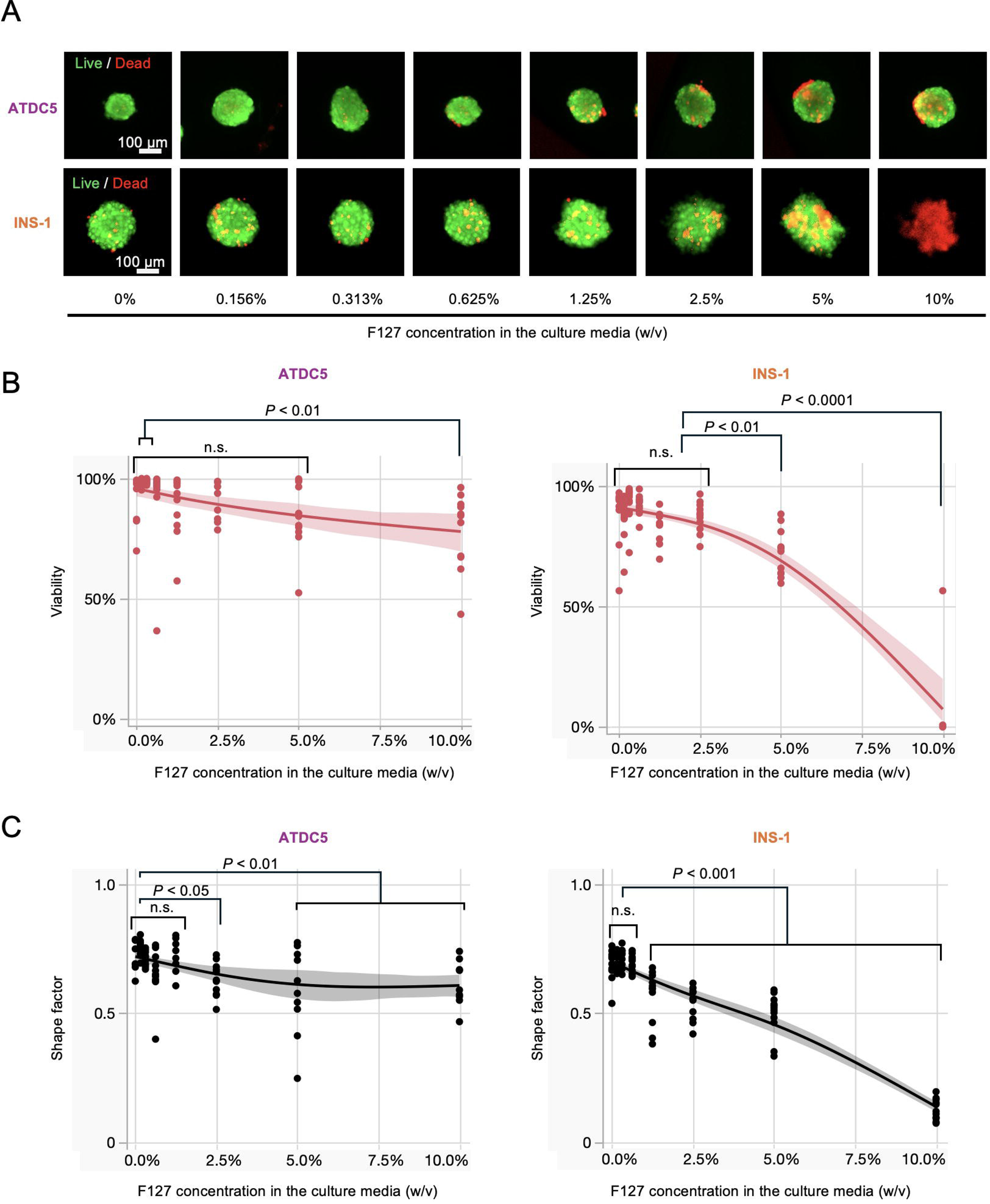
Cytotoxicity and morphological assessment of INS-1 spheroids exposed to Pluronic F127. (A) Representative fluorescence images of preformed ATDC5 and INS-1 spheroids after 24 hour exposure to Pluronic F127 added to the culture medium at 0%, 0.156%, 0.313%, 0.625%, 1.25%, 2.5%, 5%, or 10% (w/v). Spheroids were stained with fluorescein diacetate (FDA; green, live cells) and propidium iodide (PI; red, dead cells). Scale bar = 100 µm. (B) Quantification of spheroid viability based on FDA- and PI-positive areas for ATDC5 and INS-1 spheroids. The solid lines represent smoothing spline fits (λ = 3.938), and the shaded areas represent the confidence intervals of the fitted curves. (C) Quantification of spheroid shape factor for ATDC5 and INS-1 spheroids. The solid lines represent smoothing spline fits (λ = 3.241), and the shaded areas represent the confidence intervals of the fitted curves. Individual points represent individual spheroids. n = 9–12 per concentration. Group differences were evaluated by one-way ANOVA with post hoc multiple comparisons (Tukey-Kramer HSD). n.s., not significant.

### Spheroid formation kinetics on commercial non-cell-adhesive microwell plates

Before evaluating our in-house F127-coated microwells, which represented the final experimental goal of this study, we first determined the incubation time required for stable spheroid formation using commercially available non-cell-adhesive microwell plates. We seeded ATDC5 and INS-1 cells into the microwells and monitored spheroid assembly for up to 72 h (**Fig. 4A**). Representative images showed that both cell types progressively aggregated within individual microwells and formed compact spheroids within 24 h after seeding (**Fig. 4B**). Quantitative shape-factor analysis revealed significant time-dependent changes in spheroid morphology across both cell lines (one-way ANOVA, P < 0.0001 for each cell line; **Fig. 4C**). Specifically, the shape factor increased significantly from initial values at 0 hours (0.31–0.39) to stable plateaus at 24 hours (0.81–0.84; *p* < 0.0001). Beyond 24 hours, spheroid morphology remained highly stable, showing no significant differences amount the 24-, 48-, and 72-hour time points (*p* > 0.05; marked as n.s.). These kinetics demonstrate that both cell lines undergo rapid morphological condensation within the first 24 hours on non-cell-adhesive microwell plates, characterized by marked increases in sphericity, boundary smoothing, and dense cell–cell aggregation. Therefore, we selected 24 h after seeding as the time point for evaluating spheroid formation in the subsequent experiments. **Supplementary Table 4** presents detailed statistical analysis.

**Figure 4.**
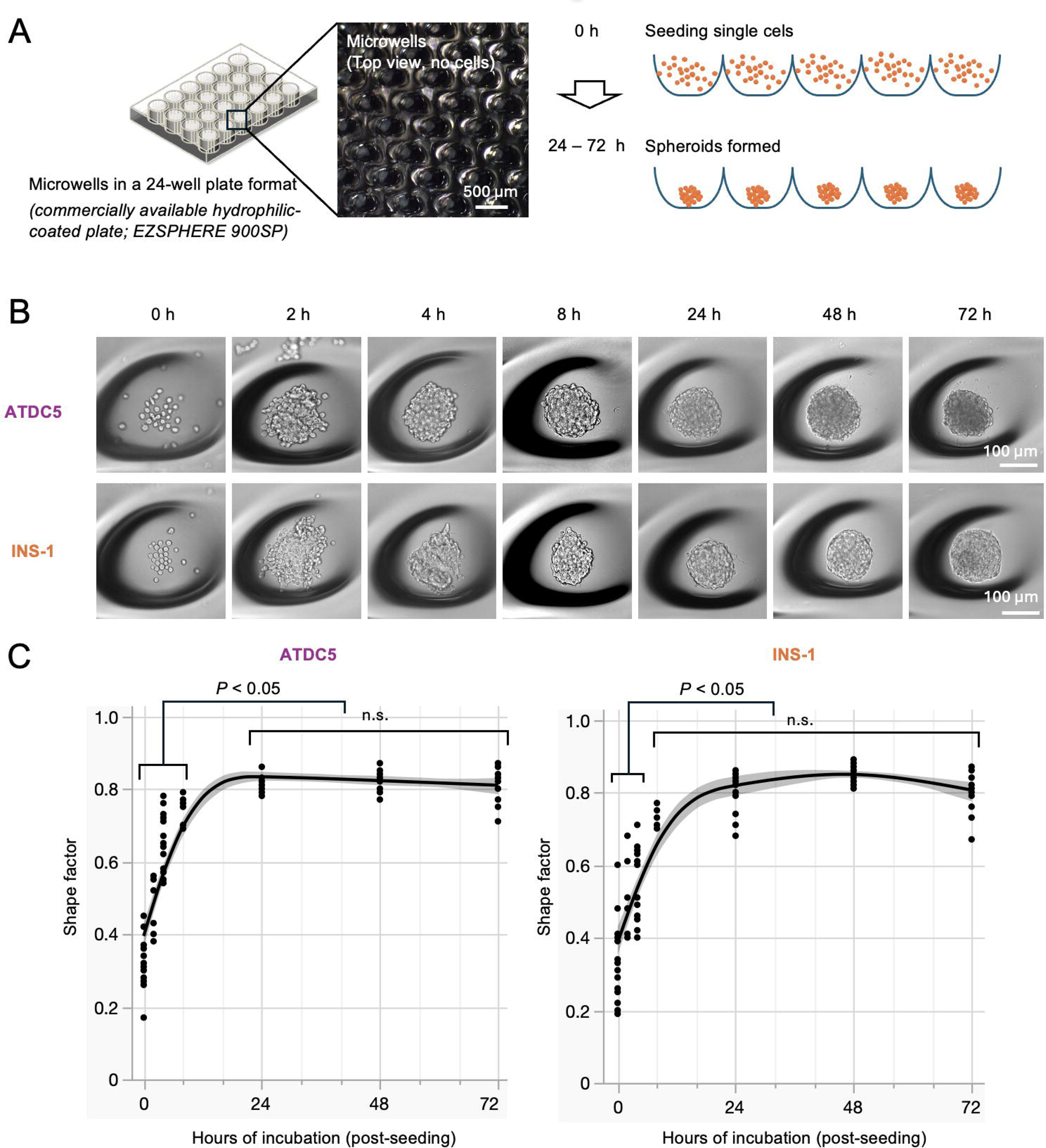
Time-dependent spheroid formation of cells in non-cell-adhesive microwells. (A) Schematic of single-cell seeding and spheroid formation in hydrophilic-coated EZSPHERE microwells. (B) Representative brightfield images of ATDC5 and INS-1 cells at 0, 2, 4, 8, 24, 48, and 72 hours after seeding into commercially available non-cell-adhesive microwell plates. Scale bars = 100 µm. (C) Quantification of spheroid shape factor over 72 hours for ATDC5 and INS-1 cells. The solid lines represent smoothing spline fits (λ = 0.179), and the gray shaded areas indicate the confidence intervals of the fitted curves. Individual points represent individual spheroids. n = 8–16 spheroids per time point. Group differences were evaluated by one-way ANOVA with post hoc multiple comparisons (Tukey-Kramer HSD). n.s., not significant.

### Enhanced spheroid formation by F127-coated microwell plates

To determine the optimal Pluronic F127 coating density for spheroid formation, we prepared microwell plates with different F127 coating densities using commercially available microwell plates after stripping their original surface coating (**Fig. 5A**). We then seeded ATDC5 and INS-1 cells into microwells coated with 0–0.0893 mg/cm^2^ Pluronic F127 and evaluated spheroid morphology after 24 h using shape factor and spheroid diameter. Representative fluorescence images demonstrated that Pluronic F127 surface coating enhanced spheroid formation in both cell lines compared with uncoated control microwells (**Fig. 5B**). Quantitative image analysis revealed that all tested F127 coating densities significantly increased the shape factor of ATDC5 aggregates relative to uncoated controls (*p* < 0.0001; **Fig. 5C**). For INS-1 cells, 0.023 mg/cm^2^ produced the highest shape factor, significantly outperforming both uncoated controls and all other coating conditions (p < 0.0001; **Fig. 5B**). Spheroid diameter analysis further highlighted distinct aggregation behaviors between the two cell lines (**Fig. 5C**). Upon F127 coating, ATDC5 cells condensed into tighter, significantly smaller spheroids. Conversely, INS-1 cells—which formed small, fragmented clusters on uncoated surfaces—consolidated into larger, well-integrated single spheroids per microwell, resulting in increased spheroid size in F127-coated microwells. Despite these cell-line-specific differences in spheroid size, F127 coating consistently improved spheroid formation across both cell lines, with coating densities between 0.011–0.045 mg/cm^2^ supporting robust, highly spherical 3D spheroid assembly. Detailed distributions of individual spheroid shape factor and diameter are shown in **Supplementary Figure 1**, and full pairwise statistical comparisons are provided in **Supplementary Table 5**.

**Figure 5.**
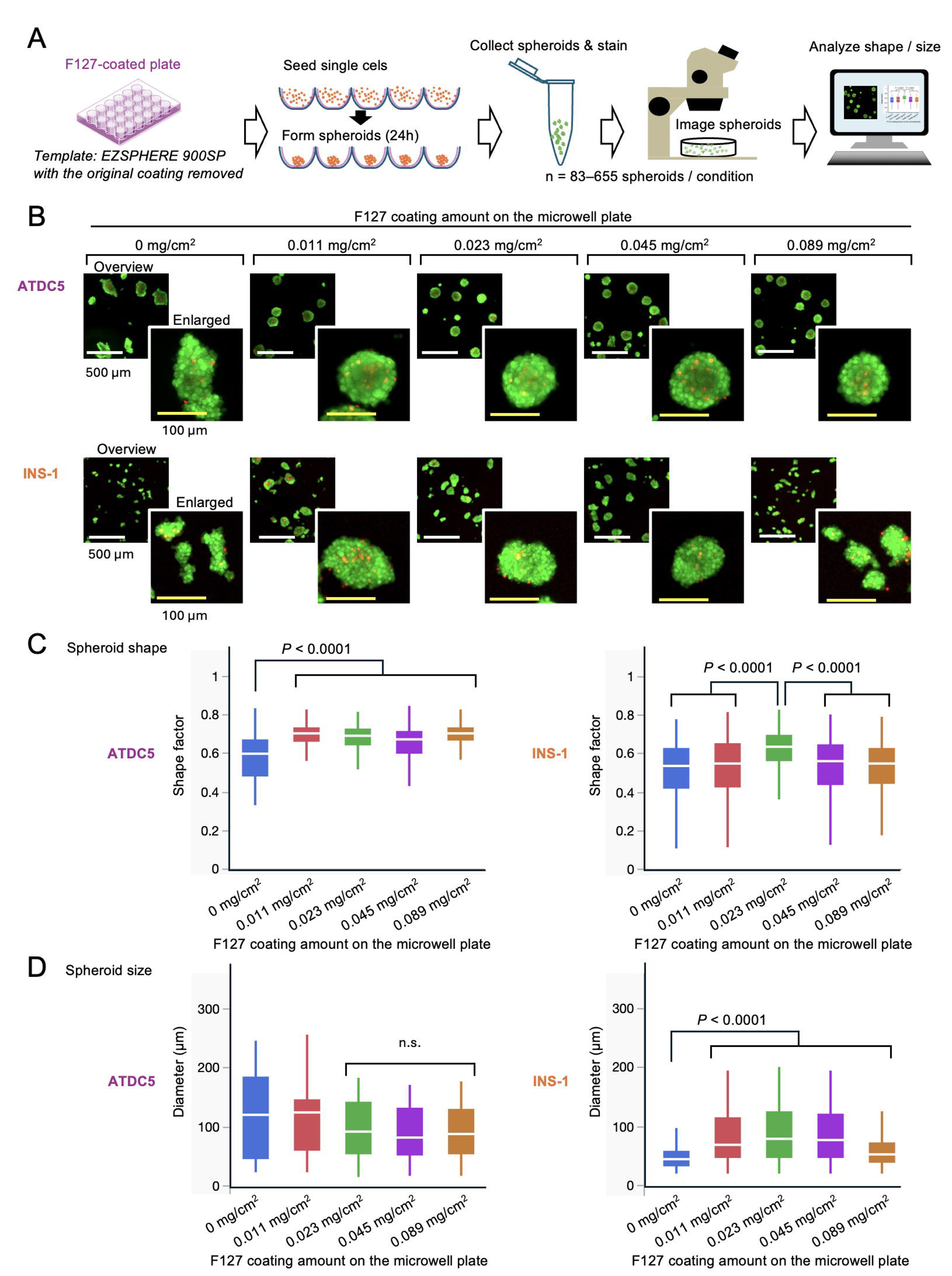
Morphological optimization of ATDC5 and INS-1 spheroid formation on Pluronic F127-coated microwells. (A) Representative fluorescence images of ATDC5 and INS-1 aggregates 24 hours after seeding into microwells coated with Pluronic F127 at 0, 0.011, 0.023, 0.045, or 0.089 mg/cm². Cells were stained with FDA and PI. Scale bars = 500 µm for overview images and 100 µm for magnified insets shown in the lower-right corner. (B) Quantification of spheroid shape factor for ATDC5 and INS-1 cells across the indicated F127 coating amounts. (C) Quantification of spheroid diameter for the same experimental groups. Box plots show the distribution of individual spheroid measurements, with the white line within each box indicating the median. n = 83–198 per F127 coating condition for ATDC5 and n = 396–655 per condition for INS-1. Group differences were evaluated by one-way ANOVA with post hoc multiple comparisons (Tukey- Kramer HSD). P < 0.0001 and n.s., not significant, are indicated in the figure.

## Discussion

Our results demonstrate that simple physical modification of polystyrene microwells using Pluronic F127 provides a robust, reproducible platform for 3D spheroid self-assembly across biologically distinct cell types. By ethanol-stripping commercial low-attachment microwell plates and re-coating them with defined amounts of Pluronic F127, we successfully reconditioned used culture plates to eliminate unwanted cell–substrate adhesion and promote uniform spheroid assembly. The reproducibility of this effect in both chondrogenic ATDC5 cells and pancreatic β-cell INS-1 cells underscores the broad generalizability of our approach. Crucially, we identified a low F127 coating density range (0.11–0.045 mg/cm^2^) that consistently supported spherical, well-compacted 3D aggregates while remaining well below cytotoxic levels. Overall, this method establishes a simple, cost-effective, and low-waste approach for generating 3D spheroids, serving as an accessible alternative to single-use culture platforms in 3D bio fabrication research.

Our surface characterization measurements provide a clear mechanistic foundation for the enhanced spheroid formation observed in F127-coated microwells. The amphiphilic structure of Pluronic F127 enables its hydrophobic poly(propylene oxide) (PPO) domain to adsorb onto the polystyrene surface, while its hydrophilic poly(ethylene oxide) (PEO) chains extend into the aqeous medium to form a hydrated interface that supresses non-specific protein adsorption ^8,12–14^. Notably, protein adsorption remained consistently low across all tested F127 coating densities, (0.011 to 0.089 mg/cm^2^), indicating that relatively low surface densities are sufficient to establish a functional non-fouling hydration layer. By dramatically reducing non-specific protein binding and cell–substrate interactions, the F127 coating creates an energetically favorable environment for cadherin-mediated cell-cell aggregation and subsequent 3D spheroid compaction. Cell-to-cell communication is critical for maintaining cell viability in 3D structures. Our cytotoxicity assays established that Pluronic F127 is a highly biocompatible coating agent preserving both cell viability and structural morphology across a wide concentration window, consistent with prior reports on its compatibility for biomedical applications^10,15,16^. We selected ATDC5 and INS-1 cells because they represent biologically distinct models with contrasting adhesion profiles and microenvironmental sensitivities; in particular, pancreatic β-cell INS-1 cells are highly sensitive to microenvironmental stress and surface-adhesion cues ^17^. Importantly, the F127 coating amounts required for optimal spheroid formation correspond to soluble concentrations far below the threshold of cytotoxicity. Based on our surface hydrophilicity and protein adsorption analyses, the maximum coating density used (0.0893 mg/cm^2^) yields surface area of a standard well in a 24-well platform. Even under the conservative assumption that the entire coating dissolves completely into 1 mL of culture medium, the resulting concentration would be only 0.25 mg/mL or 0.025% (w/v). This concentration is more than an order of magnitude lower than the levels where any cytotoxicity or morphological disruption was detected (> 1.25% w/v), confirming a broad safety margin even for sensitive cell types. Because our coating strategy relies on physical adsorption rather than covalent immobilization, long-term coating retention in physiological media warrants consideration. Covalent surface modification strategies have been investigated to prevent surfactant desorption during repeated washing or extended culture ^12^. However, persistent, long-term coating stability may not be strictly required for rapid spheroid self-assembly. In our experiments, the primary morphological transition from individual dispersed cells into compact, integrated spheroids occurred rapidly within the initial 8–24 hours. Once cells consolidate into 3D aggregates, their contact area with the underlying microwell surface decreases dramatically, diminishing the necessity for sustained, long-term surface passivation.

Interestingly, recent studies have demonstrated that supplementing culture media directly with trace concentrations of F127 can similarly induce uniform spheroid aggregation in untreated round-bottom plates, reinforcing the idea that permanent F127 surface immobilization is unnecessary for initial 3D compaction and short term culture studies^18^. Nevertheless, certain applications demand extended microwell culture, such as the long-term maintenance of pancreatic islet spheroids^19^. For these applications, future work should characterize the dissolution kinetics of physically adsorbed F127 and evaluate secondary stabilization techniques to maintain anti-adhesive properties over multi-week culture periods.

The selection of INS-1 and ATDC5 cells highlights the practical utility of this platform across distinct biological contexts. INS-1 cells serve as a validated pancreatic β-cell model relevant to pseudoislet architecture, glucose-stimulated insulin secretion, and diabetes research ^20^, whereas ATDC5 cells provide a robust chondrogenic model capable of forming dense cartilaginous matrix constructs for tissue engineering applications^21,22^. Achieving uniform spheroid formation in both models demonstrates that F127 coating effectively overcomes baseline differences in cell-type adhesion. However, a limitation of this study is that performance was evaluated using only two immortalized cell lines. Broader validation using primary cells, stem cells, and multi- cellular co-culture models will be necessary to establish fully generalizable benchmarks. Pluronic F127 provides a simple, highly reproducible approach for reconditioning polystyrene microwells to support uniform 3D spheroid formation. Rather than replacing existing commercial low-attachment platforms, this strategy offers a complementary option that expands the accessibility and sustainability of microwell-based 3D cell culture. Re-functionalizing used microwell surfaces with a widely available, non-toxic polymer is particularly advantageous for research laboratories seeking customizable or reusable culture formats ^2,18^. More broadly, this platform provides an accessible foundation for applying 3D spheroid models to high-throughput drug screening, disease modeling, tissue engineering, and regenerative medicine. By establishing a microenvironment that better recapitulates key cell-cell interactions than conventional 2D cultures, this approach enhances the physiological relevance of in vitro models and helps bridge the gap between in vitro discoveries and in vivo outcomes.

## Materials & Methods

### Maintenance and expansion of ATDC5 and INS-1 cultures

Murine chondrogenic ATDC5 (RIKEN Cell Bank, Tsukuba, Japan, Cat. # RCB0565; provided by the Skeletal Biology and Biomechanics Core of the Core Center for Musculoskeletal Biology and Medicine, UCSF, San Francisco, CA, USA) and rat insulinoma INS-1 cell lines (MilliporeSigma, Burlington, MA, USA, Cat. # SCC207) were maintained as monolayer cultures in 10 cm tissue culture-treated dishes at 37°C, 5% CO_2,_ 21% O_2_. ATDC5 growth medium contained a 1:1 (v/v) mixture of Dulbecco’s modified Eagle medium (DMEM; low glucose, with pyruvate; Gibco, Grand Island, NY, USA, Cat. # 11885084) and Ham’s F-12 (Gibco, Cat. # 11765054), supplemented with 5% (v/v) heat-inactivated fetal bovine serum (FBS; Cytiva, Marlborough, MA, USA, Cat. # SH3039603), 0.1% (v/v) penicillin-streptomycin-glutamine (100 ×; Gibco, Cat. # 10378016), and 0.1% (v/v) insulin-transferrin-selenium (100 ×; ITS-G, Gibco, Cat. # 41400045). INS-1 growth medium containing Roswell Park Memorial Institute (RPMI) 1640 (glutamine-free; Gibco, Cat. # 21870092) was supplemented with 10 mM 4-(2- hydroxyethyl)-1-piperazineethanesulfonic acid (HEPES) (Gibco, Cat. # 15630080), 50 µM β-mercaptoethanol (Sigma-Aldrich, St. Louis, MO, USA, Cat. # M3148), 1% (v/v) penicillin-streptomycin-glutamine (100 ×; Gibco), 1% (v/v) sodium pyruvate (100 ×; Gibco, Cat. # 11360- 070), and 10% (v/v) heat-inactivated FBS (Cytiva).

### Spheroid inoculation and culture parameters

Harvested ATDC5 and INS-1 cells were counted via hemocytometer and inoculated into precoated 24-well microwell arrays at seeding densities of 0.4 × 10^6^ cells/well and 0.5 × 10^6^ cells/well, respectively, in a final working volume of 1 mL media per well. Plates were cultured undisturbed at 37°C and 5% CO_2_ for 24 hours to facilitate gravity-driven self-assembly into 3D spheroids prior to downstream analysis.

### Microwell array preparation

Commercial 24-well microwell arrays (EZSPHERE 900-SP, Cat. # AG4820-900SP, Asahi Glass Company / Reprocell; ∼470 microwells/well, nominal diameter: 500 µm) served as the substrate platform. Assuming a hemispherical geometry for each microwell, the total effective surface area was estimated at 2.8 cm^2^/well (compared to a 1.9 cm^2^/well projected flat-bottom area). This 2.8 cm^2^ value was used to normalize Pluronic F127 (MilliporeSigma, Burlington, MA, USA, Cat. # P2443) coating density (mg/cm^2^). To isolate the specific effects of the F127 coating, the manufacturer’s proprietary hydrophilic surface layer was removed by treating each well with 500 µL 100% ethanol for 1 hour at room temperature (22°C). Wells were rinsed three times with sterile water and air-dried under inverted lids in a UV-C biosafety cabinet. Stripping efficiency was verified via BSA-FITC protein adsorption assay.

### Pluronic F127 coating deposition and solidification

A 10% (w/v) stock solution of Pluronic F127 in sterile water was diluted to working concentrations of 10%, 1%, 0.5%, 0.25%, 0.125%, and 0% (w/v). Stripped plates as described in **Microwell array preparation** were filled with 500 µL of coating solution and centrifuged at 300 × *g* for 5 min to eliminate trapped air bubbles. A residual volume of 25 µL was left in each well after aspirating 475 µL. Plates were dried on a rocking platform for 24 hours at 22°C, desiccated overnight at 50°C, and UV-C sterilized prior to cell seeding.

### Contact angle and surface hydrophilicity measurements

To evaluate changes in substrate hydrophilicity resulting from Pluronic F127 coating, static water contact angle measurements were performed. Flat, non-tissue culture-treated 60 mm polystyrene dishes were used as representative flat substrates to mimic the basal surface chemistry of the microwell arrays. Substrates were coated with 25 µL films of Pluronic F127 across the studied concentration range, including lower concentrations to capture the onset of hydrophilic transition. Coated dishes were air-dried with covered lids at 22°C for 24 h, followed by incubation at 50°C for 24 hours to ensure complete desiccation. To enable an unobstructed lateral view of the droplet profile, the outer side walls of the dishes were mechanically removed prior to measurement. A 10 µL droplet of sterile water was deposited onto the coated surface, and sessile drop profiles were immediately imaged using a digital single-lens reflex (DSLR) camera (EOS 1100D; Canon, Tokyo, Japan) mounted on an adjustable column (Deben Group Industries Ltd., Woodbridge, UK). Droplets were illuminated from behind using a directional light source to produce high-contrast silhouette profiles of the liquid-solid interface (shutter speed 1/15 s, f/5.6, International Organization for Standardization (ISO) 100).

### Non-fouling surface characterization

To evaluate surface non-fouling properties, 500 µL of 0.10 mg/mL BSA-FITC in PBS (pH 7.2– 7.4) was added to F127-coated 24-well plates (prepared per **Pluronic F127 coating deposition and solidification**). Uncoated wells (0% F127) served as controls. Plates were incubated at 37°C for 1 hour under light-restricted conditions. The solution was aspirated, and wells were washed three times with 1 mL PBS (rocking for 30 s per wash). After adding 500 µL fresh PBS per well, fluorescence intensity was measured using a SpectraMax iD3 microplate reader (Molecular Devices, San Jose, CA, USA) in top-read mode (exposure/emission = 485/535 nm) with default integration time, a fixed gain across all experimental groups, and plate shaking turned off.

### Kinetic time-lapse profiling of cellular self-assembly

To evaluate the kinetics of aggregate self-assembly and determine whether a 24 hour incubation window yields stable spheroid morphology, dynamic time-lapse profiling was conducted over 72 hours (n = 8–24 spheroids per condition). ATDC5 and INS-1 cells were seeded into 24-well microwell arrays with their original manufacturer coatings intact at initial densities of 0.4 × 10^6^ cells/well and 0.5 × 10^6^ cells/well, respectively. Cells were cultured under standard static conditions. To prevent fluid agitation and physical displacement of forming aggregates during plate movement, culture medium was added to fill each well to a positive meniscus. Arrays were sealed with UV-sterilized Parafilm perforated with a sterile needle to permit gas exchange while controlling headspace evaporation. Plates were incubated at 37 °C. Phase-contrast and fluorescence images were acquired at 0, 2, 4, 6, 8, 24, 48, and 72 hours using an inverted microscope (IX71; Olympus Corp., Tokyo, Japan) equipped with a 10 × objective (Numerical aperture (NA) 0.30), an IX2-SL slider, and a 4K HDMI digital camera (HD408U; AmScope, Irvine, CA, USA). Spheroid boundaries and morphological metrics were quantified as described in **Spheroid morphology and circularity analysis**.

### Spheroid morphology and circularity analysis

To quantify the geometric uniformity and aggregation kinetics of self-assembled spheroids in reused microwell arrays, brightfield images were analyzed using cellSens software (Olympus Soft Imaging Solutions, Münster, Germany). Boundary circularity was calculated using the standard area-to-perimeter shape factor showen in equation (1):

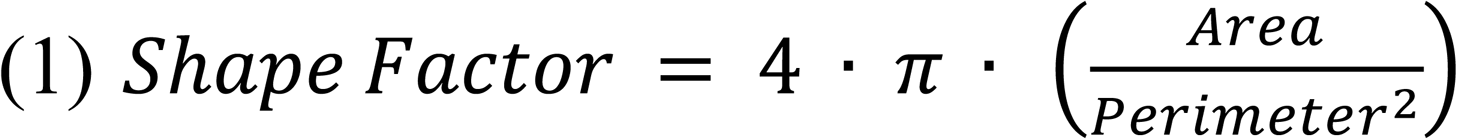

circularity value approaching 1.0 indicates a perfectly uniform circular geometry (Johnsdorf et al., 2007). Aggregate size and spatial growth dynamics were tracked over time by evaluating the effective diameter, derived from the projected surface area.

### *In vitro* biocompatibility and cytocompatibility profiling

To evaluate the biocompatibility of Pluronic F127 coatings and identify potential dose-dependent cytotoxicity, an FDA (Sigma-Aldrich)/PI (Sigma-Aldrich) cell viability assay was performed on solated 3D aggregates. Single, intact spheroids were assembled in microwell arrays as described in **Spheroid inoculation and cultivation parameters**. A 10% (w/v) stock solution was prepared by dissolving Pluronic F127 directly in complete culture medium (**Maintenance and expansion of ATDC5 and INS-1 cultures)** and serially diluted to yield eight working concentrations including 10%, 5%, 2.5%, 1.25%, 0.625%, 0.312%, 0.15%, and 0% (w/v). Individual spheroids were transferred into single wells of non-tissue culture-treated 96-well plates containing the respective F127 concentrations (n = 9–12 spheroids per condition). Plates were incubated at 37 °C, 5% CO_2_, and 21% O_2_ for 24 hours to maintain continuous chemical exposure prior to fluorometric viability imaging. To quantify the ratio of viable to non-viable cells within treated aggregates, a dual-fluorescence staining protocol using FDA and PI was implemented. A 100 µL volume containing a single spheroid and culture media was harvested directly from each well of the treatment plate and transferred into a media-preconditioned 2 mL microcentrifuge tube. For cell viability staining, a working FDA/PI solution was prepared in PBS (pH 7.4) under light-restricted conditions at 22°C by combining 5 µL of FDA stock (48 µM in acetone; Sigma-Aldrich) and 5 µL of commercial PI stock (1.5 mM) with 490 µL of PBS, yielding final working concentrations of 0.48 µM FDA and 15 µM PI ^23,24^. Spheroids were incubated in this staining cocktail for 5 min in the dark at 22°C. Following incubation, spheroids were washed with fresh PBS to remove excess dye and transferred to imaging plates for fluorescence microscopy. Fluorescence imaging was performed using a Thunder Imaging System (Leica, Wetzlar, Germany) with a 5 × objective. Whole-well tiled images were acquired to generate pan-focused projections that enabled identification of the radial live/dead interface across a two-dimensional view. Spatial distributions and signal intensities of green (viable, FDA-positive) and red (necrotic, PI-positive) fluorescence were analyzed across all Pluronic F127 concentrations and the unsupplemented control group. Digital image analysis was conducted using cellSens software (Olympus Soft Imaging Solutions, Münster, Germany). Individual spheroids were delineated to establish discrete regions of interest (ROIs), with overlapping or obscured boundaries separated manually. ROIs were filtered by minimum size thresholds to eliminate background noise, and assignments were verified to ensure a 1:1 ROI-to-spheroid ratio with consecutive indexing. For each ROI, total spheroid area and PI-positive (non-viable) area were measured. Net viable area was determined by subtracting non-viable area from total area, and percent viability was calculated as shown in equation (2):

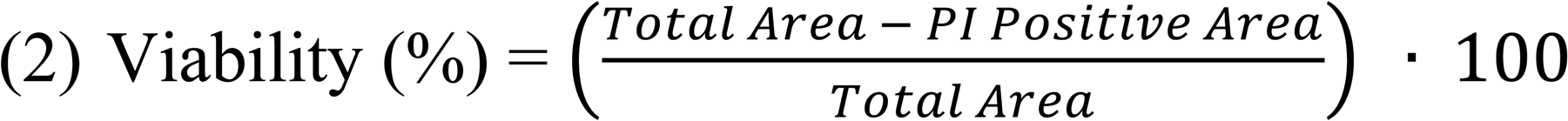

### Statistical analysis of all assays

All quantitative data visualization and statistical analyses were performed using JMP® statistical software (SAS Institute, Cary, NC, USA). Sample sizes (n) are specified in the corresponding Results sections and figure legends and represent individual spheroids or independent measurements, as indicated for each experiment. For time-course, cytotoxicity, and contact-angle datasets, data trends were visualized using JMP smoothing spline fits with confidence intervals. The smoothing parameters (λ) used for each analysis are specified in the corresponding figure legends. Spheroid shape factor and diameter distributions were visualized using box plots, with the center line indicating the median. Group differences were evaluated using one-way analysis of variance (ANOVA), followed by Tukey-Kramer honestly significant difference (HSD) tests for post hoc multiple comparisons. Statistical significance was defined as P < 0.05. P values and n.s. (not significant) are indicated in the corresponding figures, and detailed statistical results are provided in the **Supplementary Tables** where applicable.

## Acknowledgments

Portions of this work were presented at the American Association for the Advancement of Science (AAAS) 2026 Annual Meeting (Phoenix, AZ, USA, February 12–14, 2026). We thank Dr. Sunita Ho and Mandy Quan for their constructive comments and critical review of the manuscript. Abstract information is found: https://aaas.confex.com/aaas/2026/meetingapp.cgi/Paper/36111

## Author Contributions

Conceptualization: H.Komatsu. Methodology: H.Komatsu. Investigation: H.Kato; N.Ho. Formal analysis: H.Kato; N.Ho; H.Komatsu. Visualization: H.Komatsu. Writing – original draft: N.Ho; H.Komatsu. Writing – review & editing: N.Ho; H.Kato; H.Komatsu. Supervision: H.Komatsu.

## Data availability

All data generated or analyzed during this study are available from the corresponding author upon reasonable request.

## Abbreviations

BSA-FITC: Bovine serum albumin–fluorescein isothiocyanate
FBS: Fetal bovine serum
FDA: Fluorescein diacetate
HEPES: 4-(2-hydroxyethyl)-1-piperazineethanesulfonic acid
HSD: Tukey’s honestly significant difference
PBS: Phosphate-buffered saline
PEO: Poly(ethylene oxide)
PPO: Poly(propylene oxide)
PI: Propidium iodide

