## Supplemental Figure 1 for "Optimizing 3D Spheroid Formation in Microwells via a Simple Pluronic F127 Coating"

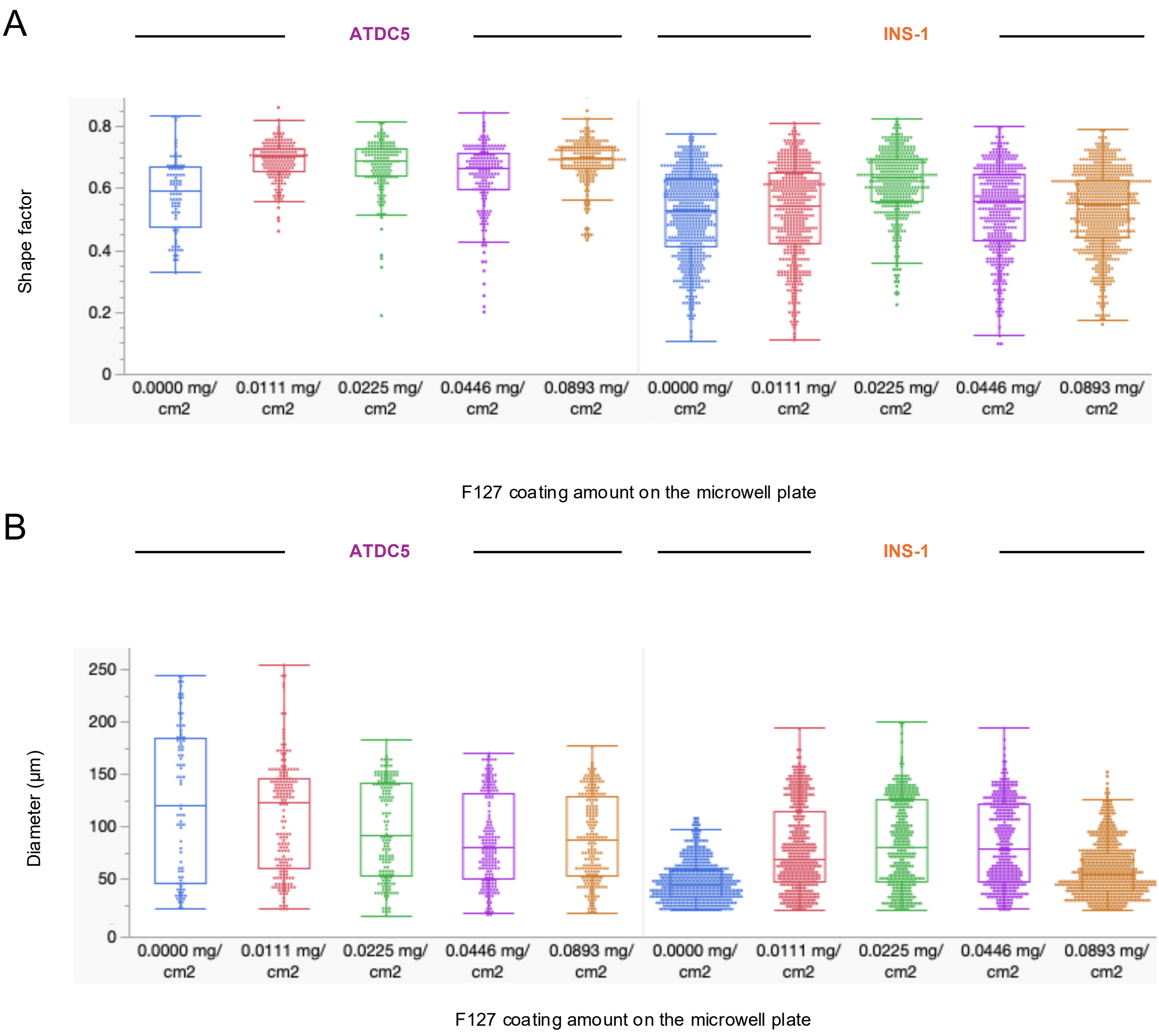


**Supplementary Figure 1. Distribution of spheroid diameter and shape factor following Pluronic F127 coating.**

(A) Distribution of spheroid shape factor for ATDC5 and INS-1 cells 24 h after seeding into microwells coated with Pluronic F127. (B) Distribution of spheroid diameter for the same experimental groups. Individual points represent individual spheroids. Box plots show the distribution of measurements, with the center line indicating the median. n = 83–198 spheroids per coating condition for ATDC5 cells and n = 396–655 per condition for INS-1 cells. Detailed statistical comparisons are provided in **Supplementary Table 5**.
